# Discovery of divided RdRp sequences and a hitherto unknown genomic complexity in fungal viruses

**DOI:** 10.1101/2020.09.08.288829

**Authors:** Yuto Chiba, Takashi Yaguchi, Syun-ichi Urayama, Daisuke Hagiwara

## Abstract

By identifying variations in viral RNA genomes, cutting-edge metagenome technology has potential to reshape current concepts about the evolution of RNA viruses. This technology, however, cannot process low-homology genomic regions properly, leaving the true diversity of RNA viruses unappreciated. To overcome this technological limitation we applied an advanced method, Fragmented and Primer-Ligated Double-stranded (ds) RNA Sequencing (FLDS), to screen RNA viruses from 155 fungal isolates, which allowed us to obtain complete viral genomes in a homology-independent manner. We created a high-quality catalog of 19 RNA viruses (12 viral species) that infect *Aspergillus* isolates. Among them, nine viruses were not detectable by the conventional methodology involving agarose gel electrophoresis of dsRNA, a hallmark of RNA virus infections. Segmented genome structures were determined in 42% of the viruses. Some RNA viruses had novel genome architectures; one contained a dual methyltransferase domain and another had a separated RNA-dependent RNA polymerase (RdRp) gene. A virus from a different fungal taxon (*Pyricularia*) had an RdRp sequence that was separated on different segments, suggesting that a divided RdRp is widely present among fungal viruses, despite the belief that all RNA viruses encode RdRp as a single gene. These findings illustrate the previously hidden diversity and evolution of RNA viruses, and prompt reconsideration of the structural plasticity of RdRp. By highlighting the limitations of conventional surveillance methods for RNA viruses, we showcase the potential of FLDS technology to broaden current knowledge about these viruses.

**Author Summary:** The development of RNA-seq technology has facilitated the discovery of RNA viruses in all types of biological samples. However, it is technically difficult to detect highly novel viruses using RNA-seq. We successfully reconstructed the genomes of multiple novel fungal RNA viruses by screening host fungi using a new technology, FLDS. Surprisingly, we identified two viral species whose RNA-dependent RNA polymerase (RdRp) proteins were separately encoded on different genome segments, overturning the commonly accepted view of the positional unity of RdRp proteins in viral genomes. This new perspective on divided RdRp proteins should hasten the discovery of viruses with unique RdRp structures that have been overlooked, and further advance current knowledge and understanding of the diversity and evolution of RNA viruses.

## Introduction

Mycoviruses, one of the most extensively studied viral groups within the virosphere, infect fungi. These viruses inhabit the insides of the rigid fungal structure, and are transmitted to other fungal cells through cell division, sporogenesis, or cell-to-cell fusion (hyphal anastomosis), and their extracellular phase is hardly ever observed (Ghabrial and Suzuki, 2009). While there are no reports of mycoviruses killing their host fungi during the fungal life cycle, some asymptomatic infections appear to be reminiscent of symbiotic relationships. Surveillance studies have revealed that mycoviruses are, to a certain extent, present in isolates of plant pathogenic fungi (Arjona-Lopez et al., 2018; Urayama et al., 2010; van Diepeningen et al., 2006). In fact, more than 200 viral species have been detected in fungi to date (Gilbert et al., 2019).

Double-stranded (ds) RNA (dsRNA) has traditionally been used as the hallmark of RNA mycovirus detection. The rapid and specific extraction method for dsRNA from fungal cells and the conventional detection method, agarose gel electrophoresis (AGE) have accelerated large-scale mycovirus screening (Morris and Dodds, 1979; Okada et al., 2015) (Arjona-Lopez et al., 2018; Khankhum et al., 2017). However, AGE cannot detect low-level mycovirus infections because its sensitivity of viral detection is relatively low.

The establishment of metagenomic and metatranscriptomic analyses for virus surveillance over the last decade has enabled more viruses to be detected in ecologically diverse environments as well as in all living creatures. These methods, which rely on deep-sequencing analysis, were expected to be more sensitive than AGE at detecting mycoviruses. Indeed, Illumina sequencing technology has successfully identified mycoviruses from fungal isolates previously considered free of these viruses by AGE (Nerva et al., 2016). Deep sequencing methods, however, are limited in their ability to identify novel viral sequences that lack homology to known virus-related sequences. Thus, finding a homology-independent method capable of detecting unidentified viruses and viral sequences in samples is required to fill this gap in viral sequence detection. Such a method would expand the list of virus-related sequences and augment current knowledge about viruses.

The multi-segmented RNA genomes possessed by some mycoviruses are coordinately replicated in the host. One segment contains an open reading frame (ORF) encoding an essential and universally seen RNA-dependent RNA polymerase (RdRp) in RNA viruses. Additional ORFs are contained in the other segment(s). In most cases, the cognate segments can be identified via the conserved sequences at their terminal ends (Urayama et al., 2018), but the conventional high-throughput sequencing methods mostly suffer from a loss of information for the terminal sequences. This technical limitation has hindered the discovery of the cognate segments and the novel viral sequences on them. To overcome this issue, we have recently developed ‘Fragmented and primer-Ligated DsRNA Sequencing (FLDS)’ technology to enable researchers to identify complete viral RNA genomes (Urayama et al., 2018; Urayama et al., 2016). This method provides reliable terminal sequences for each genome, from which the segmented RNA genomes of viruses can be determined in a homology-independent manner. This technology has higher detection power than other current methods. These advantages have allowed us to obtain highly informative complete genomes and identify uncharacterized mycoviruses using FLDS.

In the present study, we adopted FLDS technology to comprehensively screen for mycoviruses in *Aspergillus* species. Altogether, 155 isolates of *A. fumigatus* and its related species were screened, and the complete genomes of 18 viruses, as well as one with an incomplete genome, were determined for 16 fungal isolates. The FLDS-based high quality sequences we obtained supported the multi-segmented genome structure for eight viruses and uncovered novel viral sequences containing seven predicted novel ORFs in total. The most surprising finding from this screening is that one virus carries a partial RdRp lacking the essential C/D motif. The cognate segment of this virus encodes a novel ORF containing a C/D-like sequence motif. These unexpected results confirm that the ability to obtain complete, high quality viral genomes makes FLDS technology a powerful tool for expanding our current understanding of diversity in RNA viruses.

## Results

### Comprehensive screening for RNA viruses using FLDS

Altogether, 155 *Aspergillus* strains were used to screen for RNA viruses (Table S1). First, the dsRNA extracted from the mycelium of each strain was subjected to AGE. As a result, dsRNA bands were visibly detected from four *A. fumigatus* and five *A. lentulus* strains (Fig. 1). The patterns of the dsRNA bands differed from each other. The dsRNAs from the remaining 146 strains were pooled into eight groups (dsRNAs from ∼20 strains per group) and sequenced using FLDS to identify viruses that were undetectable by AGE. After RT-PCR verification, viral sequences were identified in eight strains. Overall, 17 fungi were identified that were infected with an RNA virus (Table 1), and the frequency of RNA virus-positive isolates of each species was as follows: *A. fumigatus* 8/79 (10.1%), *A. lentulus* 5/27 (18.5%) and *A. pseudoviridinutans* 4/15 (26.7%), *A. udagawae* 0/19 (0%), and *Neosartorya fischeri* 0/15 (0%) (Fig. S1). The *A. fumigatus* set, which included 20 environmental and 59 clinical isolates, contained three and five isolates that were infected with RNA viruses, respectively.

**Table 1:**
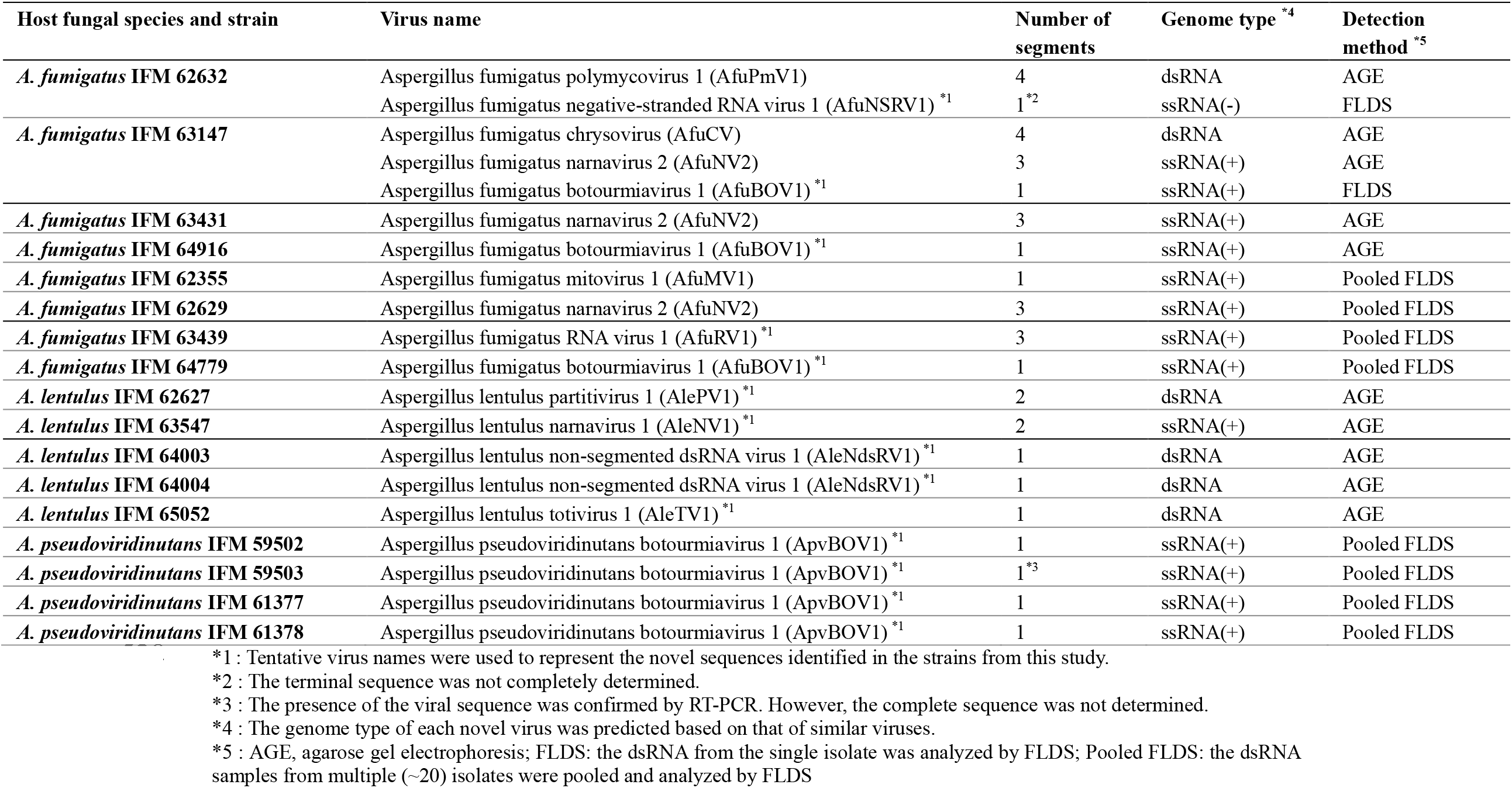
List of *Aspergillus* strains carrying viral sequences

**Fig. 1:**
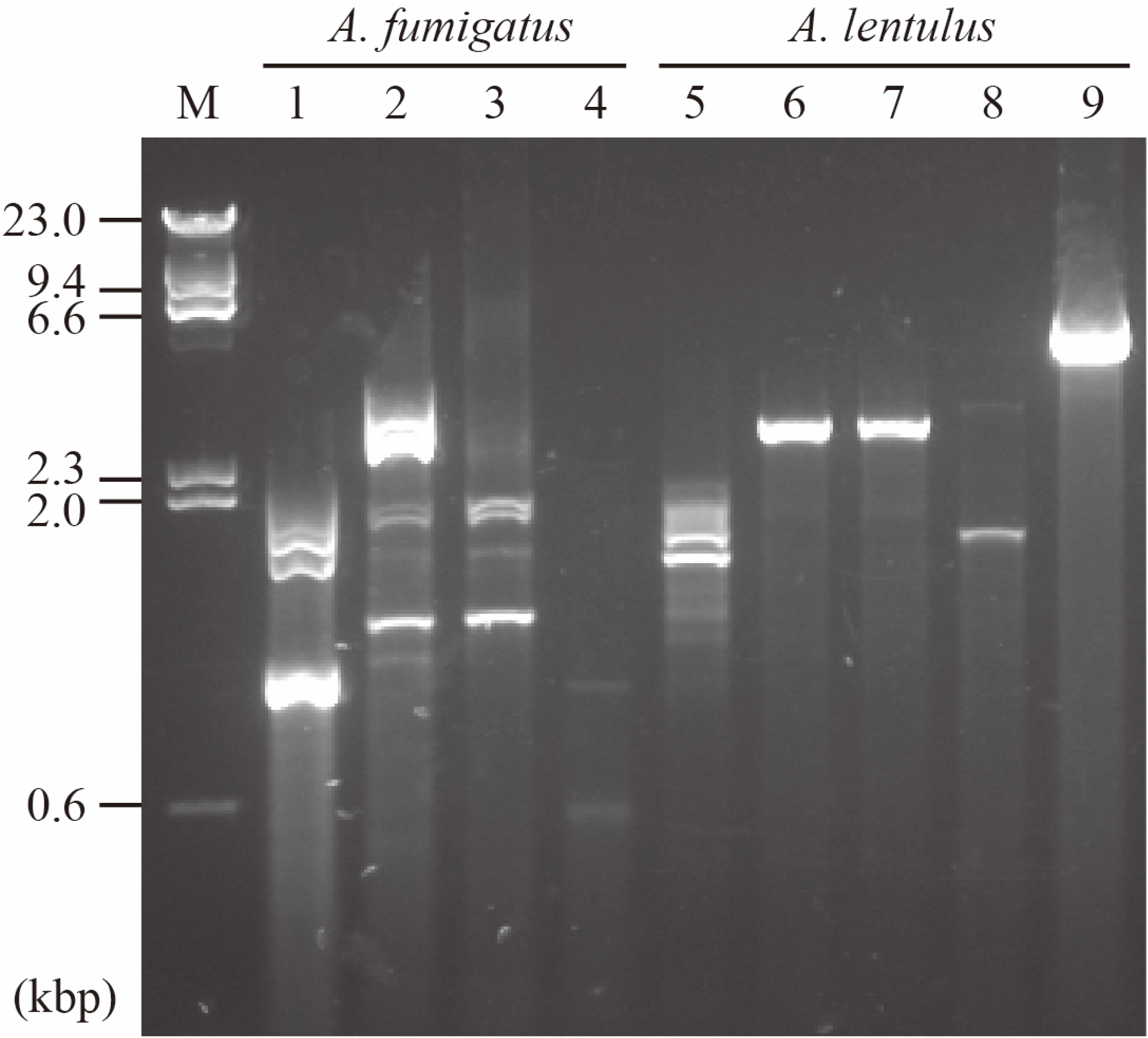
Detection of dsRNA by agarose gel electrophoresis. Comprehensive screening of 155 *Aspergillus* strains produced nine strains with positive virus-like bands. The dsRNAs from nine strains (four from *A. fumigatus* and five of *A. lentulus*) were electrophoresed, and the gel was stained with GelRed. Lanes: M, DNA marker; 1, IFM 62632; 2, IFM 63147; 3, IFM 63431; 4, IFM 64916; 5, IFM 62627; 6, IFM 64004; 7, IFM 64003; 8, IFM 63547; and 9, IFM 65052.

### Determination of segmented virus genomes

We determined the full-length viral sequences by FLDS for the strains that we identified by AGE and the pooled FLDS. Altogether, we determined the complete sequences of 16 fungal strains within three species, *A. fumigatus, A. lentulus*, and *A. pseudoviridinutans*. The complete sequence from *A. pseudoviridinutans* IFM 59503 could not be obtained despite this virus being detected by RT-PCR. The segmented viral genomes were determined according to the terminal sequence similarity among them (Fig. S2). For example, an RNA virus isolated from *A. fumigatus* IFM 62632 was found to contain four segments, whereas *A. fumigatus* IFM 63431 and IFM 62629 each contained three segments (Table 1). As a consequence, clarification of each segment resulted in us being able to differentiate the co-infecting viruses in the fungal isolates. In fact, *A. fumigatus* IFM 62632 and IFM 63147 were co-infected with two and three viruses, respectively. Altogether, 20 viruses were identified in 17 *Aspergillus* isolates, among which eight had segmented genomes (two each for bi-segments, four each for tri-segments, and two each for tetra-segments) (Table 1). The FLDS method detected the viral genomes with high sensitivity, even when multiple viruses had co-infected the same host, and it accurately discriminated segmented genomes in the viruses.

### RNA virus classification

Our BlastX analysis revealed that all 20 of the viral RNA sequences contained RdRp (Table S4). Based on the criterion that the RdRp-encoding segments sharing > 90% nucleic acid sequence identity were recognized as single operational taxonomic units (OTUs), 20 viruses fell into 12 OTUs. The virus from *A. pseudoviridinutans* IFM 59503 with a partial sequence also fell into one of the OTUs. Based on the high sequence identity (> 95%) shared with known viral sequences, four viruses were recognized as Aspergillus fumigatus polymycovirus 1 (AfuPmV1), Aspergillus fumigatus chrysovirus (AfuCV), Aspergillus fumigatus narnavirus 2 (AfuNV2), and Aspergillus fumigatus mitovirus 1 (AfuMV1). The other eight OTUs were regarded as novel viral species, and were therefore tentatively named according to the taxonomical linage of the top ‘hits’ for RNA viruses in the BlastX analysis against the NCBI nr database (Table S4). Six of eight viral species were related to dsRNA virus families (*Partitiviridae* and *Totiviridae*), positive ssRNA virus families (*Narnaviridae* and *Botourmiaviridae*), and a negative ssRNA virus family (*Betamycobunyaviridae*, a previously suggested virus family), whereas two OTUs were considered to be unclassified RNA viral linages. Notably, six of the 20 viruses identified herein, all of which had been detected by AGE, are considered to be dsRNA viruses.

*Aspergillus fumigatus* is a relatively well-studied fungal species in terms of mycovirus screening; nonetheless, three viruses, Aspergillus fumigatus botourmiavirus 1 (AfuBOV1), Aspergillus fumigatus negative-strand RNA virus 1 (AfuNSRV1), and Aspergillus fumigatus RNA virus 1 (AfuRV1), are newly identified by our FLDS-based screening. Viral isolation from related fungal species, such as *A. lentulus* and *A. pseudoviridinutans*, has never been reported; hence, the following viruses identified from the fungal species are new species: Aspergillus lentulus partitivirus 1 (AlePV1), Aspergillus lentulus non-segmented dsRNA virus 1 (AleNdsRV1), Aspergillus lentulus narnavirus 1 (AleNV1), Aspergillus lentulus totivirus 1 (AleTV1), and Aspergillus pseudoviridinutans botourmiavirus 1 (ApvBOV1).

### Structures of the novel RNA viruses identified in *Aspergillus* fungi

AfuRV1, which was identified in *A. fumigatus* IFM 63439, consists of three RNA segments (3,611, 3,447 and 1,943 nucleotides long, excluding the polyA-like region) (Fig. 2A). Each segment contains a single ORF lacking significant nucleotide identity to known sequences in the NCBI nt database. ORF1 contains methyltransferase (E-value = 1.8×10^−10^) and RdRp (E-value = 2.6×10^−82^) domains, and ORF2 contains methyltransferase (E-value = 3.3×10^−25^) and helicase (E-value = 2.5×10^−21^) domains. BlastX analysis showed that the sequence with the top hit for ORF1 was RdRp from Luckshill virus (LuV) (an unclassified ssRNA virus) (coverage, 93.0%; E-value, 0; identity, 32.7%), whereas that for ORF2 was a hypothetical protein from Cyril virus (another unclassified ssRNA virus) (coverage, 78.0%; E-value, 3.0×10^−40^; identity, 32.2%). Phylogenetic analysis of the RdRp domain showed that AfuRV1 falls into the virga-like virus clade of viruses previously isolated from invertebrates and fungi (Fig. 2B). AfuRV1, however, fell into the invertebrate-derived sub-clade containing LuV, rather than the mycovirus sub-clade. Interestingly, viruses within the virga-like clade have never been reported to possess segmented genomes. We found that the AfuRV1 genome contains an additional segment. Comparing the methyl transferase domains of ORF1 and ORF2, ORF2 fell into the virga-like virus clade, but ORF1 did not (Fig. 2C). The helicase domain in ORF2 did not fall within an established family or proposed group (Fig. 2D).

**Fig. 2:**
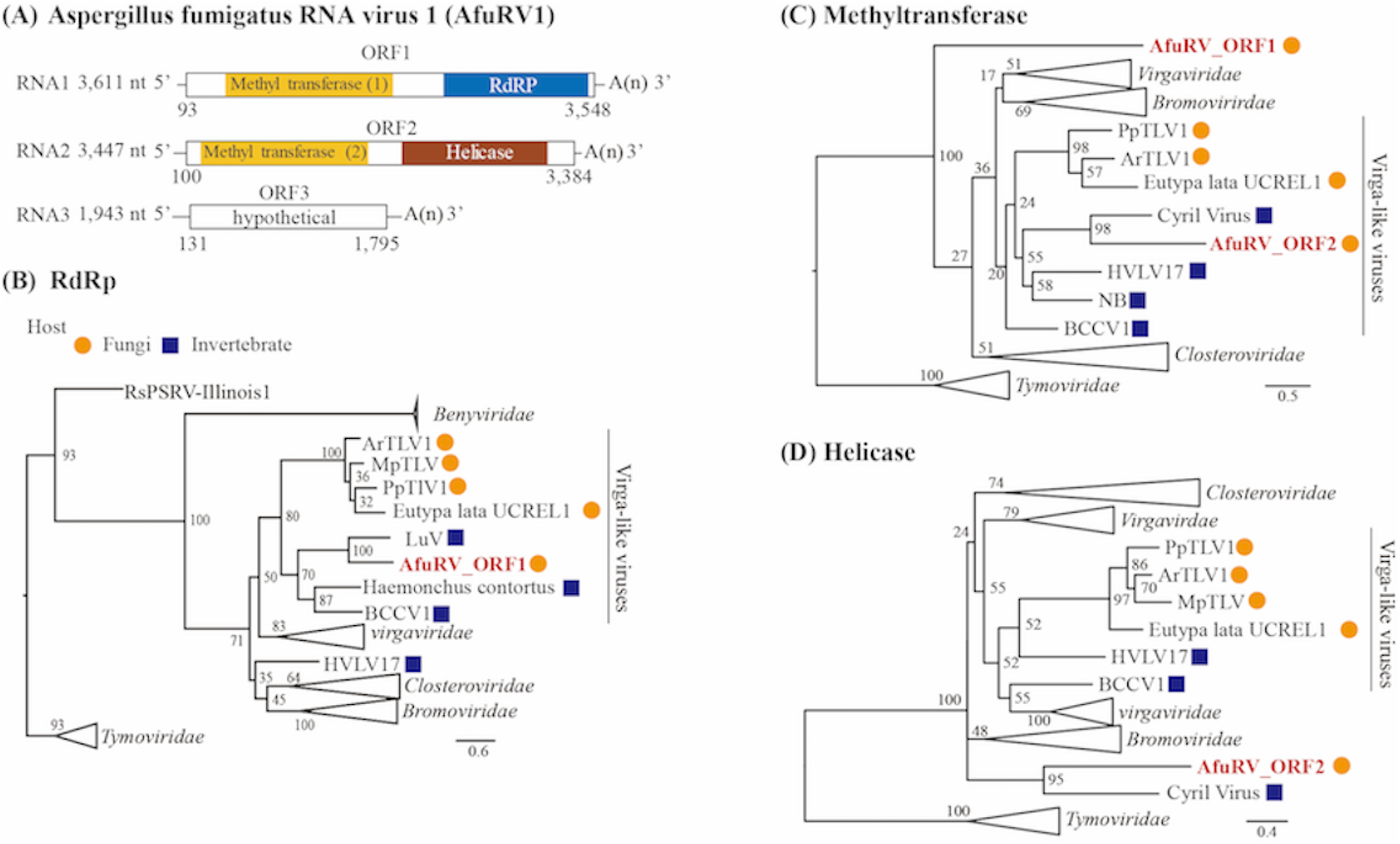
Characterization of AfuRV1. (A) The RNA genome structure model for AfuRV1. The predicted ORFs are indicated by white boxes. The domains identified as methyl transferase, RdRP, and helicase are indicated in yellow, blue and brown boxes, respectively. Molecular phylogenetic analysis on the RdRp (B), methyltransferase (C), and helicase (D) domains was performed by maximum likelihood-based methodology. The numbers indicate the percentage bootstrap support from 1,000 RAxML bootstrap replicates. The best-fitting amino acid substitution models were [LG+F+G] (B) (C) and [rtREV+F+G] (D). The accession numbers and full virus names are listed in Table S2. The scale bar represents the number of substitutions per site. The viral sequences from fungi and invertebrates are indicated by orange circles or blue squares, respectively. AfuRV1 sequences are shown in red font.

AleNV1, which originated from *A. lentulus* IFM 63547, contains two segmented genomes (3,071 and 1,814 nucleotides long, excluding the polyA-like region) (Fig. S3), whose RdRp genes share low sequence homology with that of the Beihai narna-like virus 21. The ORFs in RNA2 did not share significant similarity with known proteins. Based on its RdRp sequence, AleNV1 belongs to the well-established *Narnavirus* genus (Fig. 3). No viruses with bi-segmented genomes have been reported so far in viral genera, except LepseyNLV1 (whose complete sequence has not been reported), and MaRNAV1, which was isolated from the human trypanosomatid parasite *Leptomonas seymouri* (Grybchuk et al., 2018) and the human malaria parasite *Plasmodium vivax* (Charon et al., 2019). Thus, AleNV1 has a unique genome structure and is the first reported isolate with a bi-segmented genome among mycoviruses from the *Narnavirus* genus.

**Fig. 3:**
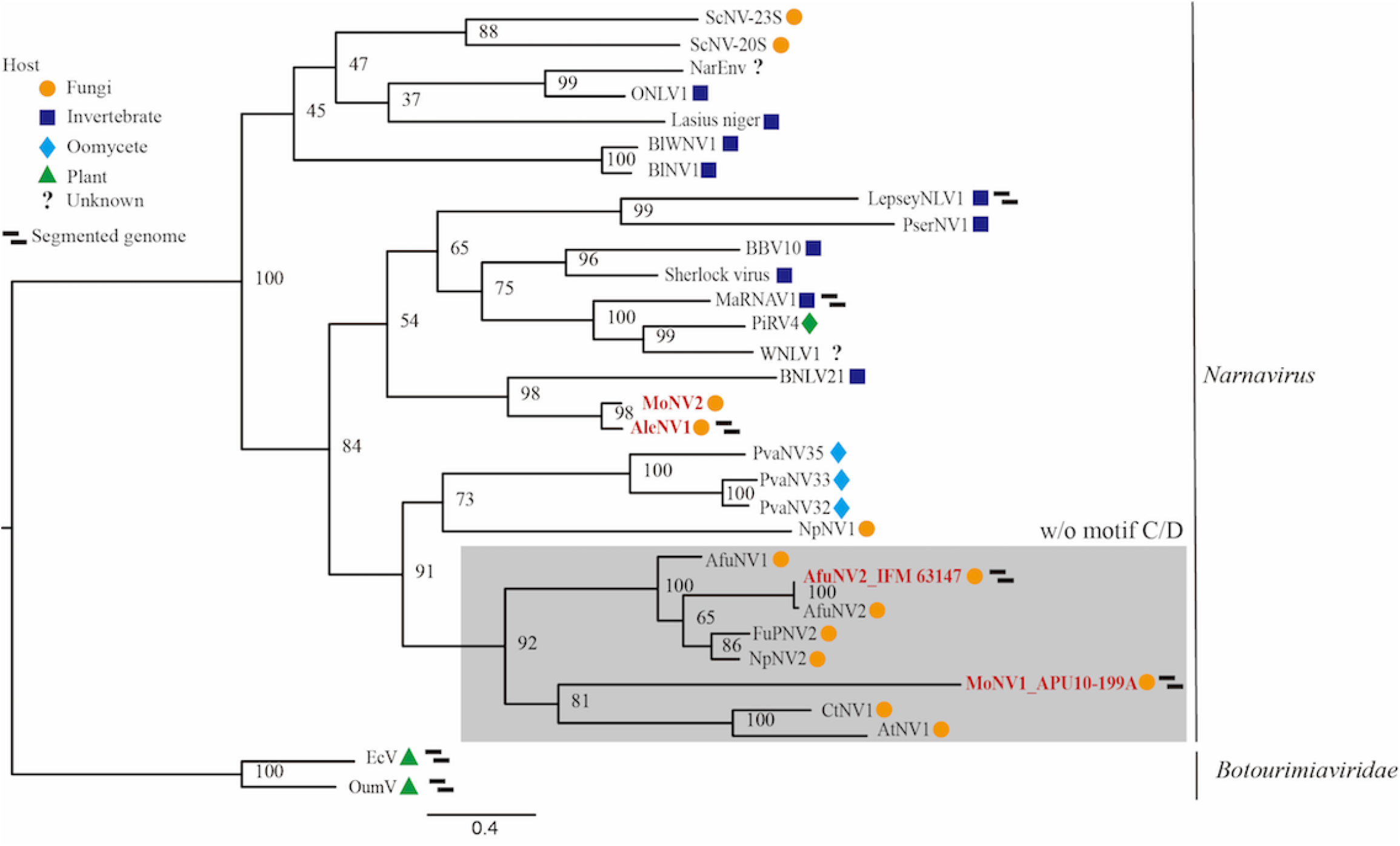
Phylogenetic tree for the RdRp from the *Narnaviridae* family. The RdRp amino acid sequences without motifs C and D from AleNV1, AfuNV2, MoNV1, and MoNV2 (without the C/D motif is shown in a gray box) and related viruses were used to construct the phylogenetic tree. The numbers indicate the percentage bootstrap support from 1,000 RAxML bootstrap replicates. The best-fitting amino acid substitution model was [LG+F+G]. The accession numbers and full virus names are listed in Table S2. The viral sequences from fungi, invertebrates, oomycetes, plants, and an unknown host are indicated by orange circles, blue squares, light blue diamonds, green triangles, and question marks, respectively. The viral sequences identified in this study are shown in red font.

### Discovery of a novel divided RdRp sequence

AfuNV2, a previously reported virus (Zoll et al., 2018), was identified in three different *A. fumigatus* isolates (IFM 63147, IFM 63431 and IFM 62629) (Table 1). Although the sequence reported by another research group is non-segmented, our FLDS-based sequencing revealed that AfuNV2 possesses RNA2 and RNA3 segments in addition to RNA1 (RdRp). This tri-segmented genome was confirmed to contain the highly conserved terminal sequences described above (Fig. S2), and was also confirmed by AGE where the band patterns were seen to correspond to the tri-segmented genome’s length (Fig. 1 lane 2 and 3). RNA2 and RNA3 are predicted to encode a large single ORF (ORF2) and multiple short ORFs (ORF3, ORF4, and ORF5) (Fig. 4A). BlastP analysis showed that the sequence with the top hit for ORF2 was RdRp from *Plasmopara viticola*-associated narnavirus 33 (PvaNV33) (coverage, 74.0%; E-value, 3E-17; identity, 23.9%), whereas ORF3, ORF4, and ORF5 share no significant similarities (e-value ≤ 1×10^−5^) with known protein sequences in the NCBI nr database or in the Pfam domain database.

**Fig. 4:**
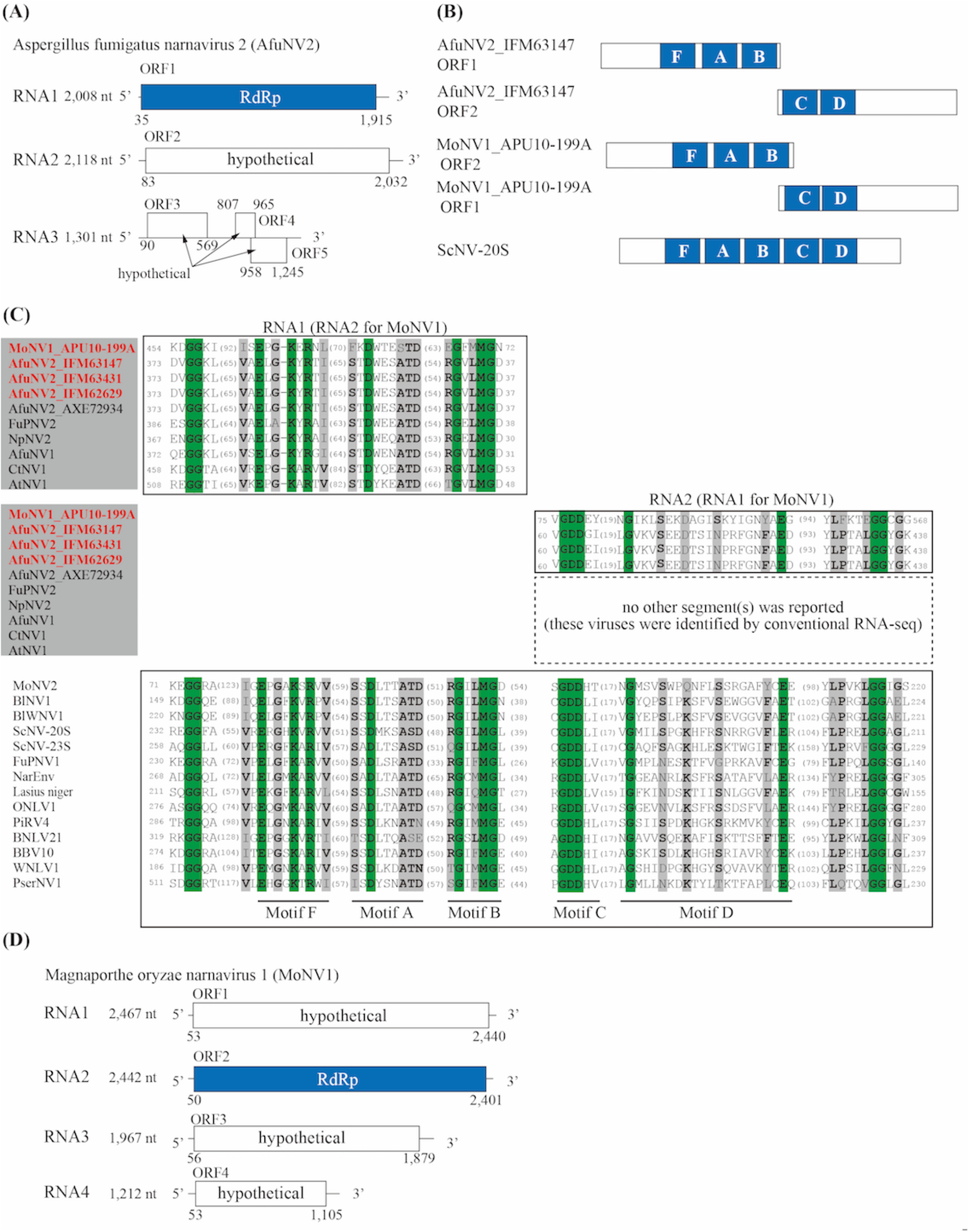
Characterization of AfuNV2 and MoNV1. (A) RNA genome structure model for AfuNV2. The predicted ORFs are represented by boxes, and the first identified RdRp domain is shown by a blue box. (B) Schematic model of RdRp protein with conserved motifs in AfuNV2, MoNV1 and ScNV-20S genomes. Conserved A–D and F motifs in RdRp from ScNV-20S, a type strain of narnavirus, are shown (Venkataraman et al., 2018, Jia, H & Gong, P, 2019). (C) Multiple alignment of the deduced amino acid sequences of the RdRp motifs for the *Narnavirus* genus. Among the 18 to 24 sequences, the amino acid positions with 100% matches and >50% matches are depicted by green and gray shading, respectively. Dominant amino acid residues at positions with >50% amino acid matches are shown in bold. The number of amino acids not shown in the alignment is noted in each sequence. The accession numbers and full virus names are listed in Table S2. (D) RNA genome structure model for MoNV1. The predicted ORFs are represented by boxes, and the first identified RdRp domain is shown by a blue box.

When the RdRp amino acid sequences were aligned, we noticed that the RdRp-encoding ORF1 from AfuNV2 lacked motifs C and D, but the well-conserved motifs (F, A and B) among the narnaviruses from other fungi were present (Fig. 4B, 4C). Interestingly, closely related narnaviruses (CtNV1, AtNV1, NpNV2, FuPNV2 and AfuNV1) were also found to lack the C and D motifs (Fig. 4C) (Lin et al., 2020). In motif C, GDD, a conserved amino acid sequence, is believed to play an essential catalytic role. Therefore, we searched for the GDD sequence in the other RNA2 and RNA3 ORFs. After much searching, we found a potential sequence in the N-terminal region of the RNA2 ORF. Sequence alignments suggested that this region includes amino acid residues that are conserved in the C and D motif regions of the RdRp proteins from narnaviruses (Fig. 4C). This was observed in three of the sequences from the AfuNV2 viruses we isolated. However, we were unable to confirm whether the aforementioned closely related CtNV1, AtNV1, NpNV2, FuPNV2 and AfuNV1 species contain RNA2 with an ORF containing a C/D-like motif sequence, because their sequences were deposited as non-segmented genomes.

To support this finding, we searched for other viruses that lack the C and D motifs in RdRp. We eventually found one known RNA virus by FLDS sequencing, Magnaporthe oryzae narnavirus 1 (MoNV1) (MN480844) from *Pyricularia* (*Magnaporthe*) *oryzae* APU10-199A, with a complete genome and reliable terminal sequences in each segment. While a non-segmented genome was reported for MoNV1, the MoNV1 we isolated has four genomic segments, but motifs C and D are missing from RdRp on RNA2 (Fig. 4D). The C/D-like motif lies within the N-terminal region of the ORF of RNA1, as was the case of AfuNV2 RNA2 (Fig. 4B). The ORFs on RNA3 and RNA4 from MoNV1 contain no predicted proteins or domains with significant homology. In addition to MoNV1, another narnavirus, MoNV2, co-infected the same strain. MoNV2 has a non-segmented genome containing a single ORF that shares significant nucleotide identity with RdRp from AleNV1 (coverage, 99.0%; E-value, 0; identity, 74.61%) (Fig. 3). The RdRp from MoNV2 harbors a C/D motif (Fig. 4C). Collectively, the results for these two different fungal species (*A. fumigatus* and *P. oryzae*) suggest that certain narnaviruses carry RdRp sequences that are divided into two different ORFs, with one ORF containing motifs F, A and B, and the other potentially containing motifs C and D.

## Discussion

DsRNA mycoviruses infect a wide range of fungal species including Ascomycota, Basidiomycota, Glomeromycota, and Mucoromycotina (Gilbert et al., 2019, Turina et al. 2018, Kartali et al. 2019). Some viruses can affect growth, development, toxin production, and pathogenicity in their fungal hosts. Fungal viruses have not been intensively studied despite their potential impact on ecology, agriculture, food security, and public health. In this study, we evaluated the recently developed FLDS technology for its sensitivity and accuracy of RNA viral sequence detection. Several new viral sequences were successfully identified as complete genomes. A most intriguing finding was the presence of a divided RdRp gene. No examples like this irregular form have been reported in any other RNA viruses known to infect living creatures. This result could be only accomplished by FLDS-based sequencing because of its ability to produce high-quality end-to-end sequences. This demonstrates its utility for capturing previously unidentified viral sequences from RNA viruses.

Large-scale AGE screening for RNA viruses was conducted on a set of more than 300 *Aspergillus* strains (Bhatti et al., 2012), and 6.6% of the detected strains contained dsRNA. Elsewhere, of the 86 *A. fumigatus* clinical isolates that were examined 18.6% contained dsRNA (Refos et al., 2013). Here, we detected dsRNA by AGE from 9/155 strains (5.8%) of *Aspergillus* species. Another advanced screening method, metatranscriptomics, was recently used to detect viral sequences, and 72 assembled virus-related sequences from 275 isolates of plant pathogenic fungi were identified (Marzano et al., 2016). The searches were based on identifying sequences with significant identity to known amino acid-encoding viral sequences. Notably, only 15% of the sequences were predicted to be derived from dsRNA, while 73% and 12% were predicted to be derived from positive-sense RNA or negative-sense RNA viruses, respectively. More recently, a pipeline to efficiently detect viral sequences from a transcriptomics dataset was proposed (Gilbert et al., 2019). This was also based on sequence similarity to RdRp. Of the 569 RNA-Seq samples, 59 complete mycoviral genomes were identified in 47 datasets, 34 viruses (57%) were predicted to be dsRNA viruses, and 88% were new species. Here, 10 viruses were identified by FLDS that were overlooked by AGE. All the overlooked viruses were predicted to be ssRNA viruses. We consider that the amount of dsRNA recovered as a replicative intermediate of ssRNA is low. Nonetheless, FLDS captured sequences derived from ssRNA viruses. Therefore, the highly sensitive, high-throughput sequence-based screening of FLDS is a powerful tool for constructing deep and wide viral catalogs.

One apparent limitation occurs with homology-based viral sequence detection whereby the novel ORFs accumulated by knowledge-based updating can overlook those lacking homology to known virus-related ORFs, resulting in a biased list of viral sequences. Our FLDS-based screening was clearly able to circumvent this issue by identifying six viral species with segmented genomes and discovering seven ORFs that had never been recognized as virus-related sequences (ORFs encoding hypothetical proteins with unknown functions, Table S4).

The discovery of segmented genomes provides evolutionary insight into the origin of certain mycovirus groups. AfuRV1 has three segmented genomes and contains two methyltransferase domains in different segments. As far as we are aware, RNA viruses with multiple methyltransferase domains have never been reported in published literature. Phylogenetic analysis revealed that the methyltransferase domain of ORF2 (MT2) and the RdRp domain of ORF1 belong to a virga-like virus clade. In contrast, the methyltransferase domain of ORF1 (MT1) and the helicase domain of ORF2 fall into an unclassified group (not the virga-like virus clade). This suggests that the ancestor of AfuRV1 acquired MT1 and its helicase domain from a different virus. Identifying the origins of these domains and the evolutional history of AfuRV1 is not straightforward because MT1 and the helicase domain are distantly related to the domains of known viruses. By FLDS analysis, we identified segmented genomes in AfuNV2 and AleNV1 that both belong to the *Narnaviridae* family. Interestingly, segmented genomes have only been reported for LepseyNLV1 (a protozoal virus) and *Botourmiaviridae* family plant viruses, and they have not been reported among the *Narnaviridae* family or closely-related ssRNA viruses (Grybchuk et al., 2018; Rastgou et al., 2009). Hence, this is the first identification of *Narnaviridae* family mycoviruses with multi-segmented genomes. It is noteworthy that AfuNV2 and AleNV1 were not classified as belonging to the sub-clade that includes LepseyNLV1. Furthermore, AfuNV2 and AleNV1 narnaviruses are not closely related to each other and have different genome structures. This suggests that during evolution of the *Narnaviridae* family and its relative ssRNA viruses, the acquisition of segments and changes in genome structure occurred independently in each host kingdom. We cannot, however, rule out the possibility that non-RdRp encoding segments are satellite RNAs that tentatively coexist. Further investigations in this area are required.

The gene encoding RdRp is universally present among RNA viruses (Wolf et al., 2018), and all known RNA viruses encode RdRp in a single ORF (King et al., 2012). Unexpectedly, we found that viruses in a certain clade of *Narnaviridae* encode an RdRp that lacks catalytic domains C and D, and a different coexisting ORF encodes the missing domains. This irregular genome structure was identified in two different viruses from two different fungal hosts, *Aspergillus* and *Pyricularia*. Besides these two viruses, this group includes other viruses isolated from *Fusarium* (FuPNV2), *Neofusicoccum* (NpNV2), *Cladosporium* (CtNV1), and *Alternaria* (AtNV1) (Fig. 3). According to the deposited sequences, these viruses appear to lack domains C and D of RdRp. These sequences were not obtained by FLDS; thus the corresponding ORF with a GDD motif or cognate genomes were unlikely to be detected; indeed, no such sequences have been deposited. The deposited sequence dataset from other narnaviruses supports the possibility that imperfect RdRp proteins exist in a certain group of mycoviruses that infect a wide range of the Ascomycetes phylum. As described above, the ORF2 of AfuNV2 showed low but certain similarity to the RdRp sequence from PvaNV33. Interestingly, PvaNV33 showed certain similarities not only to ORF2 but also to ORF1 of AfuNV2 (Fig. S4). Moreover, sequences of RdRp from PvaNV32 (GenBank: QIR30311.1) and PvaNV35 (GenBank: QIR30314.1) that formed a single clade with PvaNV33 also shared certain similarities to both ORFs of AfuNV2. This complementary genomic structure suggests that the RdRps from these *Plasmopara viticola*-associated narnaviruses are the molecular ancestors from which ORF1 and ORF2 of AfuNV2 were derived by division. During the preparation of this manuscript, another research group reported on the identification of Magnaporthe oryzae narnavirus virus 1 (MoNV1) from an *M. oryzae* strain isolated in China (Lin et al. 2020). The viral sequence was deposited as a single genome with RdRp lacking domains C and D, which concords with our results for the MoNV1 we identified in our work. This consistency in the viral sequences from different countries suggests that viruses with irregular genome structures are widely distributed, meaning that the event leading to the atypical structures did not occur locally. Our finding of divided RdRp sequences raises questions about how divided RdRps function in host cells and whether atypical structures are limited to mycoviruses. Answering these questions is of great interest to us.

In conclusion, our FLDS-based screening for mycoviruses using 155 fungal isolates has led to the discovery of novel species and novel segmented genomes in some of the viruses. Some viral genomes have novel structures that would not have been captured by conventional methods. Thus, FLDS-based screening has potential to tap into unexplored diversity in RNA viruses.

## Materials & Methods

### Strains and culture conditions

The *Aspergillus* strains used in this study are listed in Table S1. These strains were cultured in potato dextrose broth (PDB) with reciprocal shaking (120 rpm) for up to 5 days at 30°C or 37°C. All the strains were provided by the National BioResource Project (https://nbrp.jp/). *Pyricularia* (*Magnaporthe*) *oryzae* APU10-199A (Higashiura et al., 2019) was cultured in PDB for 1 week at 30°C before harvesting.

### RNA extraction

Fungal mats (fresh weight, 100 mg) were disrupted in liquid nitrogen in a mortar or using FastPrep 24 (MP Biomedicals Inc., OH, USA). dsRNA and ssRNA purification was performed as described previously (Urayama et al., 2020; Urayama et al., 2018; Urayama et al., 2016). In brief, total nucleic acids were manually extracted from the ground cells with sodium dodecyl sulfate–phenol. dsRNA was purified using the cellulose resin chromatography method (Morris and Dodds, 1979; Okada et al., 2015) and subjected to AGE analysis. To obtain sequence-grade dsRNA, the remaining DNA and ssRNA were removed with amplification grade DNase I (Invitrogen, Carlsbad, CA, USA) and S1 nuclease (Invitrogen). Total RNA was extracted from the pulverized samples using the TRIzol Plus RNA Purification Kit (Invitrogen), and the eluted RNA was treated with amplification grade DNase I (Invitrogen) and purified using RNA Clean & Concentrator-5 (Zymo research, Irvine, CA, USA).

### Sample pooling, sequence library construction and sequencing

When dsRNA band(s) were visible by AGE, the dsRNA samples from each isolate were individually prepared for viral genome sequencing by FLDS. When no visible dsRNA band or bands were observed, the dsRNA samples from up to 20 isolates were pooled into a single sample (referred to as pool 1–8), following FLDS analysis (referred to as pooled-FLDS analysis).

dsRNA was converted into dscDNA by the FLDS method (Urayama et al., 2018). Each purified dsRNA was fragmented by ultrasound using a Covaris S220 ultrasonicator (Woburn, MA, USA), and a U2 adapter was ligated to each dsRNA fragment using T4 RNA ligase (Takara Bio Inc., Kusatsu, Japan). After denaturing the product, single-stranded (ss) cDNA (sscDNA) was synthesized using the SMARTer RACE 5′/3′ Kit (Takara) with a U2-complementary primer. dscDNA was obtained by PCR with a U2-complementary primer and a universal primer mix (provided by the SMARTer RACE 5′/3′ Kit).

cDNA libraries were constructed as described previously (Urayama et al., 2018). Each 300 bp of the paired-end sequences from each fragment were determined on the Illumina MiSeq platform (Illumina, CA, USA).

### Data processing

Clean reads were obtained by removing low-quality, adapter and low-complexity sequences as described previously (Urayama et al., 2018). For the RNA virome analyses, contaminated rRNA reads were removed by SortMeRNA (Kopylova et al., 2012). According to a previous method (Urayama et al., 2018), the cleaned reads were subjected to *de novo* assembly using the CLC Genomics Workbench version 11.0 (CLC Bio, Aarhus, Denmark). The resulting assemblies were manually examined and extended using the assembly Tablet viewer (Milne et al., 2010). Where the terminal end of a contig ended with same bases for more than 10 reads or a polyA sequence was present, the position was recognized as the terminal end of the RNA genome. When a contig had termini at both of its ends, it was considered to be the full-length sequence of the RNA genome. Multi-segment genomes were judged according to the terminal sequence similarities of the segments. BlastN and BlastX programs (Camacho et al., 2009) were used to identify sequence similarities among known nucleotide sequences and protein sequences, respectively.

### Phylogenetic analysis

ORF prediction was performed by ORFfinder, after which Pfam domain searching was conducted (Finn et al., 2016). Phylogenetic analysis of the helicase and methyltransferase domains of RdRp, which were based on the amino acid sequences obtained from the NCBI nr database, were aligned using MUSCLE (Edgar, 2004) in MEGA6 (Tamura et al., 2013). The accession numbers of the sequences used for the analyses are listed in Table S2. Alignment-ambiguous positions were removed with trimAl (Capella-Gutiérrez et al., 2009). Maximum likelihood-based phylogenetic analyses were performed using RAxML (Stamatakis, 2014), and bootstrap tests were conducted with 1,000 samplings. The amino acid substitution model was selected by Aminosan (Tanabe, 2011) using Akaike’s information criterion (Sugiura, 1978). To visualize phylogenetic trees, Fig-Tree (Rambaut, 2014) was used.

### Reverse transcription (RT) PCR (RT-PCR)

To detect RNA viruses from host fungi in pooled samples, RT-PCR analyses were performed using specific primer sets (Table S3) as described previously (Urayama et al., 2014). After pricking the mycelia grown on a potato dextrose agar several times with a toothpick, the toothpick was then dipped into the one-step RT-PCR reaction mix in a PCR tube and twisted three times. One-step RT-PCR was performed using the Super-Script III One-Step RT-PCR System with Platinum Taq (Invitrogen) according to the manufacturer’s protocol. To support the sequences provided by FLDS, the 3′ end sequences of RNAs 1 and 2 from Aspergillus fumigatus RNA virus 1 (AfuRV1) were confirmed by One-step RT-PCR using an oligo (dt) primer and specific primers (Table S3). Total RNA was used as the template. PCR products were run on 1% agarose gels and the visualized fragments were excised, purified using the FastGene Gel/PCR Extraction Kit (Nippon Genetics, Tokyo, Japan), and then used for direct Sanger sequencing.

### Data accessibility

Data sets supporting the results of this study are available in the GenBank database repository (Accession Nos. DDBJ: LC553675–LC553714) and the Short Read Archive database (Accession No. DDBJ: DRA010415).

## Acknowledgments

This research was supported by a grant from the Institute for Fermentation, Osaka, and in part by a Grant-in-Aid for Scientific Research (18H05368 and 20H05579) from the Ministry of Education, Culture, Sports, Science and Technology (MEXT) of Japan and a Grant-in-Aid for Scientific Research on Innovative Areas from the Ministry of Education, Culture, Science, Sports, and Technology (MEXT) of Japan (No. 16H06429, 16K21723, and 16H06437). We thank Drs. Hiromitsu Moriyama and Shin-ichi Fuji for providing *M. oryzae* APU10-199A. We thank Sandra Cheesman, PhD, from Edanz Group (https://en-author-services.edanzgroup.com/ac) for editing a draft of this manuscript.

## Figure Legends

**Fig. S1: Frequency of RNA virus-positive isolates and the detection method.** AGE: agarose gel electrophoresis.

**Fig. S2: Multiple alignments of the 5′-and 3′-terminal regions of the RNA sequences.** Nucleotide positions with 100% matches among the sequences are depicted by green shading. Numbers at the beginning and end of each sequence represent the nucleic acid positions.

**Fig. S3: Genome structure models for the viruses identified in this study.** The predicted ORFs are shown by a box. The domains identified in RdRp and the coat protein (CP) are shown by blue and gray boxes, respectively. Hypothetical proteins are shown as ‘hypothetical’.

**Fig. S4: Schematic model of RdRp protein from PvaNV33 and comparison with AfuNV2.** Conserved A–D and F motifs in RdRp from PvaNV33 and AfuNV2 are shown. The regions with similarity between two proteins are shown in a striped square.

